# Norepinephrine improves attentional orienting in a predictive context

**DOI:** 10.1101/485813

**Authors:** Amelie J. Reynaud, Mathilda Froesel, Carole Guedj, Sameh Ben Hadj Hassen, Justine Clery, Martine Meunier, Suliann Ben Hamed, Fadila Hadj-Bouziane

**Affiliations:** INSERM, U1028, CNRS UMR5292, Lyon Neuroscience Research Center, ImpAct Team, Lyon, F-69000, France; University UCBL Lyon 1, F-69000, France; CNRS, UMR5229, Institut des Sciences Cognitives Marc Jeannerod

**Keywords:** Atomoxetine, monkey, LATER, visuo-spatial attention, reaction time

## Abstract

The role of norepinephrine (NE) in visuo-spatial attention remains poorly understood. Our goal was to identify the attentional processes under the influence of NE and to characterize these influences. We tested the effects of atomoxetine injections (ATX), a NE-reuptake inhibitor that boosts the level of NE in the brain, on seven monkeys performing a saccadic cued task in which cues and distractors were used to manipulate spatial attention. We found that when the cue accurately predicted the location of the upcoming cue in 80% of the trials, ATX consistently improved attentional orienting, as measured from reaction times (RTs). These effects were best accounted for by a faster accumulation rate in the valid trials, rather than by a change in the decision threshold. By contrast, the effect of ATX on alerting and distractor interference was more mitigated. Finally, we also found that, under ATX, RTs to non-cued targets were longer when these were presented separately from cued targets. This suggests that the impact of NE on visuo-spatial attention depends on the context, such that the adaptive changes elicited by the highly informative value of the cues in the most frequent trials were accompanied by a cost in the less frequent trials.

## 1. Introduction

Visuo-spatial attention is a pervasive function that enables us to selectively process visual information through prioritization of a spatial location while setting aside other locations. It depends on the fronto-parietal network and is under the influence of several neuromodulators including dopamine (DA), acetylcholine (ACh) and norepinephrine (NE) (see Noudoost and Moore 2011). While a systematic approach to understand the role of DA and ACh in visual-spatial attention has been carried out over the years, the role of NE is currently less understood (see Noudoost and Moore 2011).

In particular, only a handful of studies have addressed the role of NE transmission in visuo-spatial attention and its sub-components (alerting, orientating and executive control; Posner 1980, Petersen and Posner 2012), and the results are inconsistent. Petersen and Posner (2012) suggest a specific role of NE in the maintenance of high sensitivity to incoming stimuli i.e. the alerting sub-component (Petersen and Posner, 2012). At least two studies provide evidence in support of this (Witte and Marrocco 1997; Coull et al. 2001). Evidence of the contribution of NE to spatial orienting is more mitigated (Clark et al. 1989; Coull et al. 2001; Witte and Marrocco 1997). As to attentional executive control, the third attentional sub-component, reaction times to identical external events have been shown to be affected by general task context, and to be much faster in highly predictive contexts than in less predictive contexts (Los 1996; Los et al. 2001; Albares et al. 2011; Wardak et al. 2012). While there is, to our knowledge, no direct evidence for an effect of NE onto this attentional sub-component, a recent study shows a selective increase in pupil size, an indirect index of NE activity, in the presence of highly predictive cues (Dragone et al., 2018). This, thus, suggests a possible interaction between NE and attentional executive control.

Here, we focused onto these three specific attentional components, namely alerting, spatial orienting and executive control and we aimed at 1) clarifying the components that are under the influence of NE and 2) characterizing the specific action of NE onto them.

We thus tested seven monkeys in a saccadic cued task derived from the attentional network task (Posner 1980). This task allows manipulating the focus of attention by using cues that precede the appearance of the target. We used a context where the cue accurately predicted the spatial location of the upcoming target in 80% of the trials. A distractor could also appear simultaneously with the target to examine the subjects’ ability to filter distractors out when planning their saccadic movement. We tested the monkeys under two pharmacological conditions: after saline administration used as the control condition and after atomoxetine administration, a NE reuptake inhibitor that increases the level of NE in the synaptic clefts. To investigate whether alerting and orienting were affected by a boost in NE transmission, we computed attentional network scores from the reaction times (Fan et al. 2002). To identify changes driven by task context and executive control, we compared RTs in highly predictive tasks and in less predictive tasks. To investigate how these attentional processes were affected by a boost in NE transmission, we used the LATER model (linear approach to threshold with ergodic rate; Carpenter and Williams 1995) to test whether changes in RT distributions following NE modulation were better accounted for by a change in signal accumulation rate, signing a perceptual process, or a change in decision threshold, signing a top-down process (Noorani and Carpenter 2016). One could expect either 1) a global non-specific NE effect onto all three attentional components; 2) an NE effect specific to the alerting non-selective attentional component or 3) an NE effect specific to the dynamic/flexible components of attention, namely orienting and executive control. Our observations speak in favor of the last prediction.

## 2. Methods

### 2.1. Subjects

Seven rhesus monkeys (*Macaca mulatta*) aged 5-14 years participated to this study, three females (monkeys CA, GU and CE) and four males (monkeys EL, TO, HN and DO). Animals had free access to water (CE, CA and GU) or food (EL, TO, HN and DO) and were maintained on a food (CE, CA and GU) or water (EL, TO, HN and DO) regulation schedule, individually optimized to maintain stable motivation and performance. This study was conducted in strict accordance with Directive 2010/63/UE of the European Parliament and the Council of 22 September 2010 on the protection of animals used for scientific purposes and approved by the French Committee on the Ethics of Experiments in Animals (C2EA) CELYNE registered at the national level as C2EA number 42.

### 2.2. Experimental set-up

Monkeys were seated in a primate chair in a sphinx position, with the head immobilized via a surgically implanted plastic MRI-compatible head post (CE, TO, EL, HN, DO) or a non-invasive head restraint helmet (CA and GU) (Hadj-Bouziane et al. 2013), in front of a computer screen (distance: 57cm for CE, CA and GU; 78cm for EL, TO, HN and DO). Gaze location was sampled at 120 Hz using an infrared pupil tracking system (ISCAN Raw Eye Movement Data Acquisition Software) interfaced with a program for stimulus delivery and experimental control (Presentation^®^).

### 2.3. Behavioral task

A testing session consisted of alternations of two types of runs: mixed runs and pure runs. In both types of runs, monkeys were required to fixate a central cross to initiate the trial. Then, the target appeared randomly in the left or right side of the screen (10 degrees of eccentricity), and monkeys had to saccade as fast as possible to the target location and hold fixation during 300ms (EL, TO, HN and DO) or 500ms (GU, CA and CE) to receive a reward (fruit juice or water). In the mixed runs, derived from the attentional network task (Posner 1980), several conditions were intermixed, while in the pure runs, only one condition was presented to the animals. For 4 monkeys (EL, TO, HN and DO), the color of the central cross changed across the type of runs (red or yellow cross for mixed and pure runs, respectively).

In the mixed runs (figure 1A), for 80 % of the trials, a peripheral cue, a white dot or a grey square, was flashed for 100ms prior to the target onset on one side of the screen, accurately predicting the upcoming target location (‘*valid* cue’). In the remaining 20% of the trials, the cue was either absent (‘*no* cue’), or presented on the opposite side of target location (*‘invalid* cue’), or two cues were simultaneously presented (*‘neutral* cue’). In addition, a distractor, a red circle or a red square, could appear simultaneously with the target onset, either in the same or in the opposite hemifield as the target (distance target-distractor: 4.5° for GU, CA and CE and between 2.1° and 3.2° for EL, TO, HN and DO). The ‘*no distractor*’, ‘*same hemifield*’ and ‘*opposite hemifield*’ conditions were intermixed and equally distributed across trials. Monkeys were required to fixate the target and ignore the distractor. In the majority of the animals (except CE), the cue-target interval (CTI) varied across trials to prevent anticipatory responses. CTIs were optimized for each monkey in order to maximize cue validity/invalidity effects, which were key in quantifying the attention orientation effects (200-300-400ms for GU and CA, 100ms for CE, 150-200-250ms for EL and TO, 200-250-300ms for HN, 140-180-240ms for DO). The pure runs did not include any cue nor any distractor. These runs served to quantify the effect of NE on task context by comparing RTs on these trials to the same trials performed in the mixed runs (i.e. taking place in a context in which cued trials were most frequent). The mixed runs included ~ 90 trials for monkeys CE, CA, GU, ~ 150 trials for monkeys EL, TO, ~ 300 trials for monkey HN and ~ 400 trials for monkey DO. Pure runs included ~ 20 trials for monkeys CA, GU, ~ 50 trials for monkeys EL, TO, ~ 100 trials for monkey HN and ~ 150 trials for monkey DO. Note that only mixed runs were presented to monkey CE.

**Figure 1.**
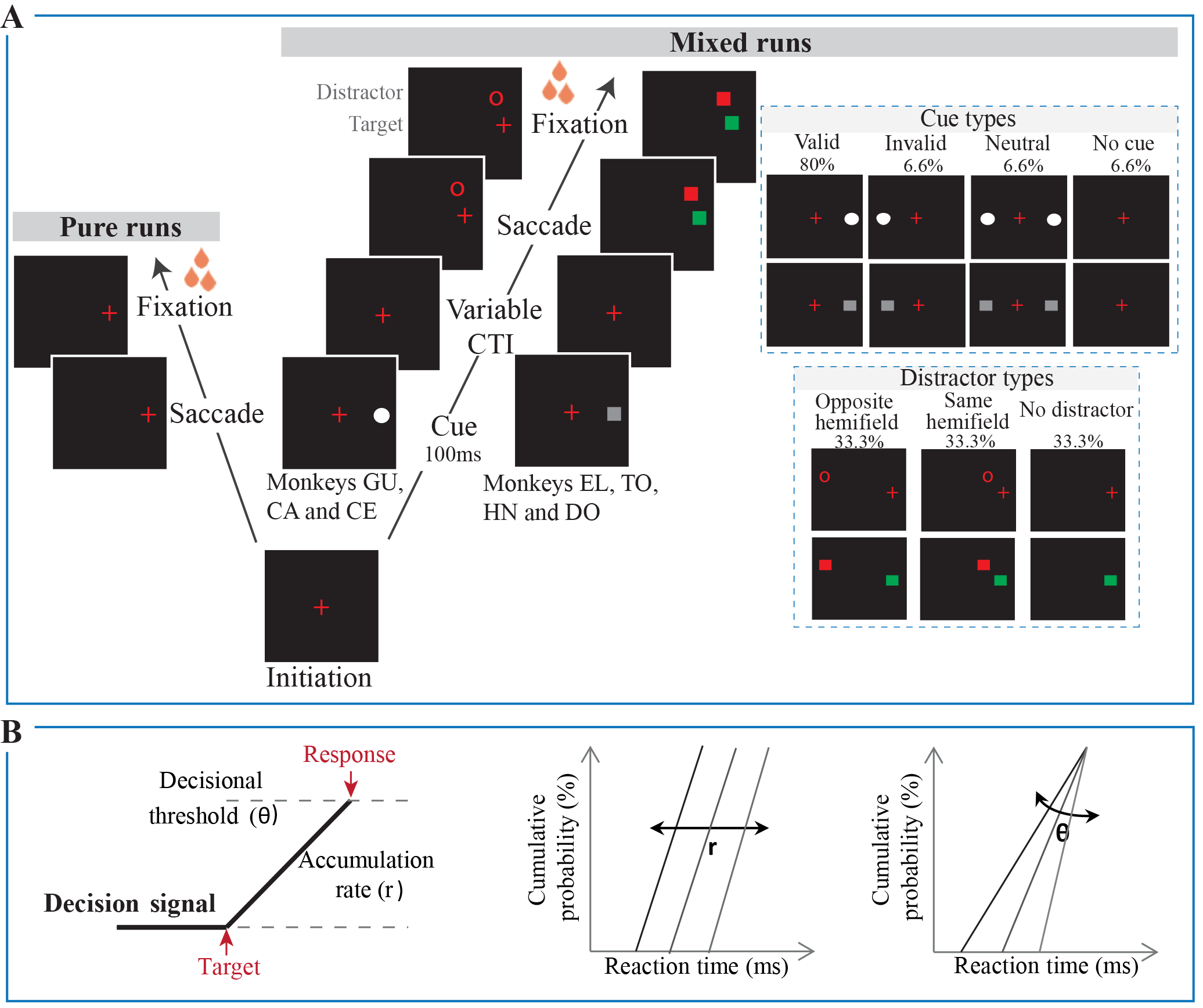
Behavioral task and LATER model. **A:** In the mixed runs (spatial attention cued task), monkeys initiated the trial by fixating the red cross. Then, a cue was flashed and the monkey was required to keep his gaze on the red cross. The cue could either be valid, invalid, neutral or absent. After a cue-target interval (CTI), the target appeared on one side of the screen. Simultaneously, a distractor could appear in the same or opposite hemifield of the target location. The monkey had to ignore the distractor and saccade to the new target location to successfully complete the trial and receive a reward. The pure runs did not include any cue nor any distractor and monkey had to saccade to the target location to successfully complete the trial and receive a reward. **B:** According to the LATER model, RT is the culmination of a decisional signal which starts at the apparition of the target, rises in response with a constant linear rate (r) and ends with the initiation of a response at the decision threshold (θ) (left panel). Cumulative RT distributions are plotted as reciprobit plots, so that each distribution corresponds to a line. On this plot, the change of accumulation rate is embodied by a shift of the lines and the change of the decisional threshold by a swivel of the lines (right panel).

The overall structure of the task was similar for all animals. Only the physical characteristics of the stimuli (cues, target and distractors) and the timings varied across animals depending on their previous experience with the task and their overt behavior (figure 1A).

### 2.4. Drug administration

Once the animals reached stable performance and were accustomed to intramuscular injections, atomoxetine, a NE reuptake inhibitor (ATX, Tocris Bioscience, Ellisville, MO) and saline (control) administration sessions began. ATX or saline was administered intramuscularly 30 min prior to testing (Gamo et al. 2010). Each experiment started with one or two weeks of saline administration, followed by 3 to 4 weeks of testing with different doses of ATX: 0,1mg/kg, 0,5mg/kg, 1mg/kg and 1,5mg/kg. For a given week, the same dose of ATX was administered every day to the animals. Note that the dose of 1,5mg/kg was tested only in the two younger monkeys (GU and CA). In total, for each animal, we collected 4 to 6 sessions with each dose of ATX and 1 to 5 sessions of saline condition.

### 2.5. Data analysis

The data were analyzed separately for each monkey. Eye movements were visually inspected with a customized toolbox implemented in MATLAB.

#### 2.5.1. Pupil diameter

We computed the averaged normalized pupil diameter, in the trial initiation period (500ms before the cue onset), for each animal and each pharmacological condition. In each trial, the mean pupil diameter across this 500ms window was divided by the root mean square separately for each animal. These measures were compared across runs and pharmacological conditions.

#### 2.5.2. Number of Trials

We examined the number of initiated trials (i.e. Figure 1A: completion of the first step: initiation) and the number of correct trials. A trial was considered correct after the animal reached and fixated the correct target location within the imparted time (270ms for DO, 300ms for EL and HN, 350ms for TO or 500ms for CA, GU and CE). Incorrect trials corresponded to either incomplete trials, anticipations (RT < 80ms), saccades with artifacts related to blink or trials where saccades were made to the wrong target location or to the distractor location.

#### 2.5.3. Reaction times (RTs)

##### 2.5.3.1. Attentional scores in mixed runs

To assess the effect of cues and distractors on RTs in mixed runs, we computed four scores derived from the attentional network scores (Fan et al. 2002) and integrating the effect of distractors (Walker and Benson 2013): alerting score, orienting score, remote distractor score and proximal distractor score. Given that these different conditions were randomly presented within runs, these scores were calculated for each run. Runs where the number of trials per cue type was under-represented (i.e. less than 3 trials) were excluded.

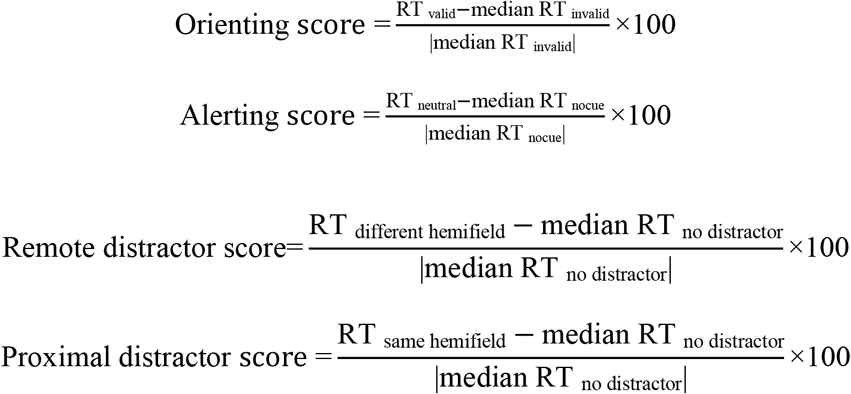

##### 2.5.3.2. RT distributions in mixed runs

We used the LATER model (linear Approach to Threshold with Ergodic Rate) to examine changes in RT distribution for each attentional process (Noorani and Carpenter 2016). This model proposes that RT is the culmination of a decisional process which starts at the onset of the target, rises in response with a constant linear rate (r) and ends with the initiation of a response at the decision threshold (θ) (figure 1B left panel). According to this model, a change in RT distribution can be explained by a change in the accumulation rate or in the decision threshold. Cumulative RT distributions are plotted as reciprobit plots, so that each distribution corresponds to a line. On this plot, the change of accumulation rate is embodied by a shift of the lines and the change of the decision threshold by a swivel between them (figure 1B right panel). To characterize how RT distribution was affected by trial type (i.e. to characterize a given attentional process or NE effect), we calculated the log likelihood ratio that the difference between one RT distribution and the other is accounted for by a shift or by a swivel. A negative log likelihood ratio represents a change in accumulation rate between the two distributions (i.e. a shift) and a positive log likelihood ratio represents a change in decisional threshold (i.e. a swivel).

##### 2.5.3.3. Non-cued trials in pure versus mixed runs

We compared RTs of correct trials in non-cued trials presented in the pure and in the mixed runs. We also used the LATER model to examine the changes in RT distribution between the types of runs.

### 2.6. Statistical analysis

We used linear mixed models (using the ‘lme4’ package for R, Bates et al. 2014) to examine the effect of ATX on the different variables computed above, for each monkey. As a first step, we defined a model containing the most appropriate random effects (i.e. factors of non-interest). Random effects were thus introduced sequentially, and their effect on model fit was assessed through Likelihood Ratio Tests (LRT): residuals of each model were compared, and the one with significantly lower deviance as assessed by a chi-squared test was chosen (table 1). We then tested the effect of pharmacological condition as fixed factor to evaluate the effect of ATX on pupil diameter, number of initiated and correct trials and the different attentional scores. To evaluate the effect of ATX on RTs in the different cue conditions, we tested the effect of pharmacological condition and cue condition as fixed factors. Finally, post-hoc comparisons were carried out using pairwise comparisons through the ‘lsmeans' package for R (p-adjusted with false discovery rate method, Lenth 2016) to assess the effect of the different doses of ATX (and the different cue conditions when assessing their effects on RTs).

**Table 1.** Linear mixed models in statistical analysis. For each variable, we defined a model containing the most appropriate random effects (i.e. factors of non-interest) and we tested the effect of fixed factor (i.e. factor of interest). Random effects were sequentially introduced, and their effect on model fit was assessed through Likelihood Ratio Tests (see methods).

|  | Variables | Family of linear mixed model | Random factors | Fixed factors |
| --- | --- | --- | --- | --- |
| Mixed run | Pupil diameter | Gaussian | Runs nested in sessions | Pharmacological condition |
|  | Number of initiated trials | Binomial | CTI; Target position; Cue condition; Distractor condition; Runs nested in sessions | Pharmacological condition |
|  | Number of correct trials | Binomial | CTI; Target position; Cue condition; Distractor condition; Runs nested in sessions | Pharmacological condition |
|  | RTs | Gaussian | Target position; Cue condition; Distractor condition; Runs nested in sessions | Pharmacological condition; Cue condition |
|  | Alerting score | Gaussian | CTI; Target position; Distractor condition; Runs nested in sessions | Pharmacological condition |
|  | Orienting score | Gaussian | CTI; Target position; Distractor condition; Runs nested in sessions | Pharmacological condition |
|  | Remote distractor score | Gaussian | CTI; Target position; Cue condition; Runs nested in sessions | Pharmacological condition |
|  | Proximal distractor score | Gaussian | CTI; Target position; Cue condition; Runs nested in sessions | Pharmacological condition |
| Pure run | Number of initiated trials | Binomial | Target position; Runs nested in sessions | Pharmacological condition |
|  | Number of correct trials | Binomial | Target position; Runs nested in sessions | Pharmacological condition |
|  | RTs | Gaussian | Target position; Runs nested in sessions | Pharmacological condition |

To determine whether a particular strategy was used between the different conditions (different cue conditions, distractor conditions or types of runs), LATER model log likelihood ratio tests were performed for each subject. To evaluate the effect of ATX on the strategy, we performed a Kruskal-Wallis non-parametric paired test on the group of subjects.

## 3. Results

In the results section below, the ATX dose-response curves are provided for pupil diameter (figure 2) and attentional orienting effect (figure 3). Other results are detailed for the highest, and most effective, dose of ATX (1.0mg/kg for monkeys CE, EL, TO, HN, DO and 1.5mg/kg for monkeys GU and CA).

**Figure 2.**
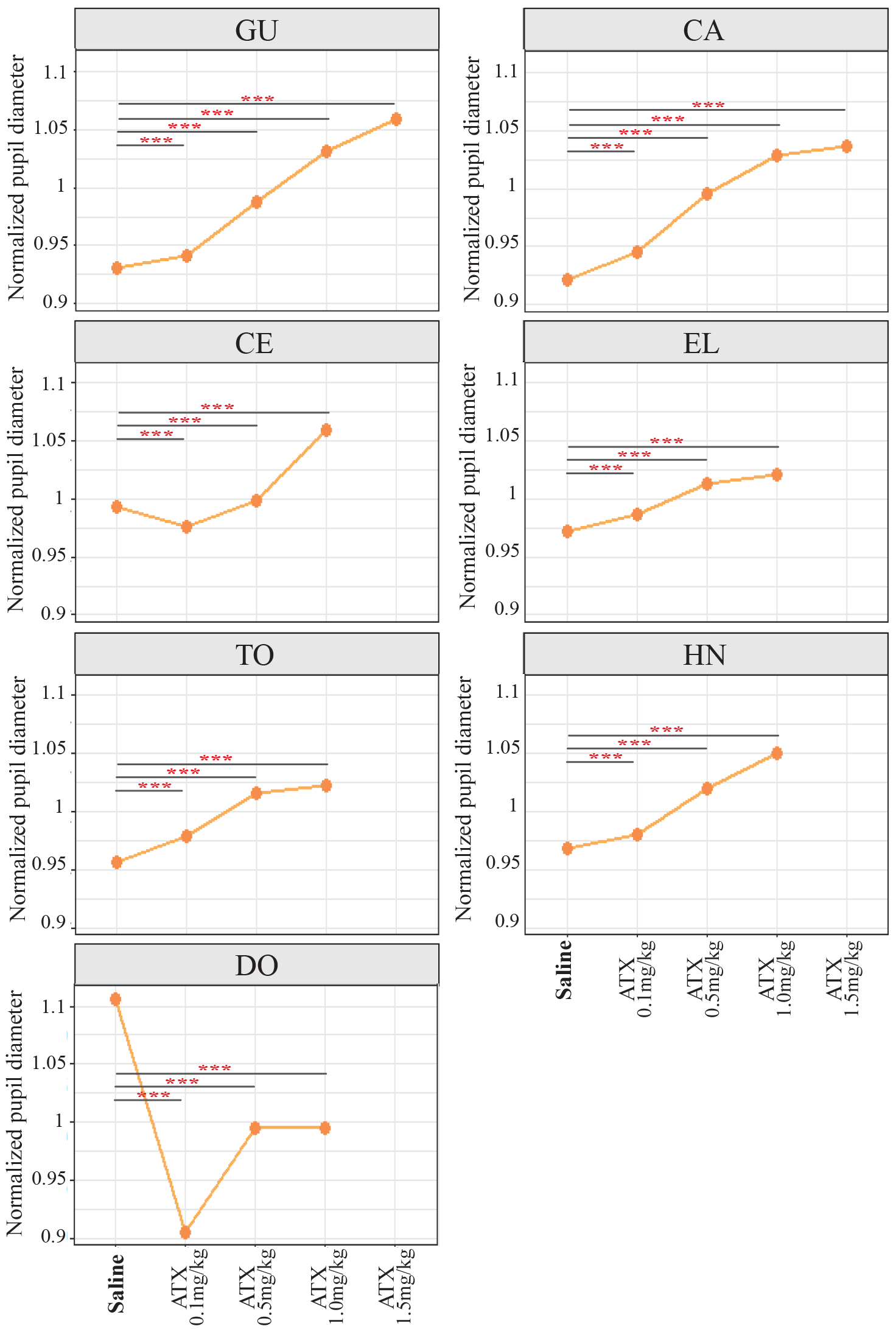
ATX effect on pupil size. For each animal and each pharmacological condition, we computed normalized averaged pupil diameter (mean±s.e) during the initiation period (fixation cross). ATX significantly increased pupil diameter as a function of the dose, in most of the monkeys, during the initiation period. \**:p-value<0.01; **:p-value<0.05; ***:p-value<0.001*.

**Figure 3.**
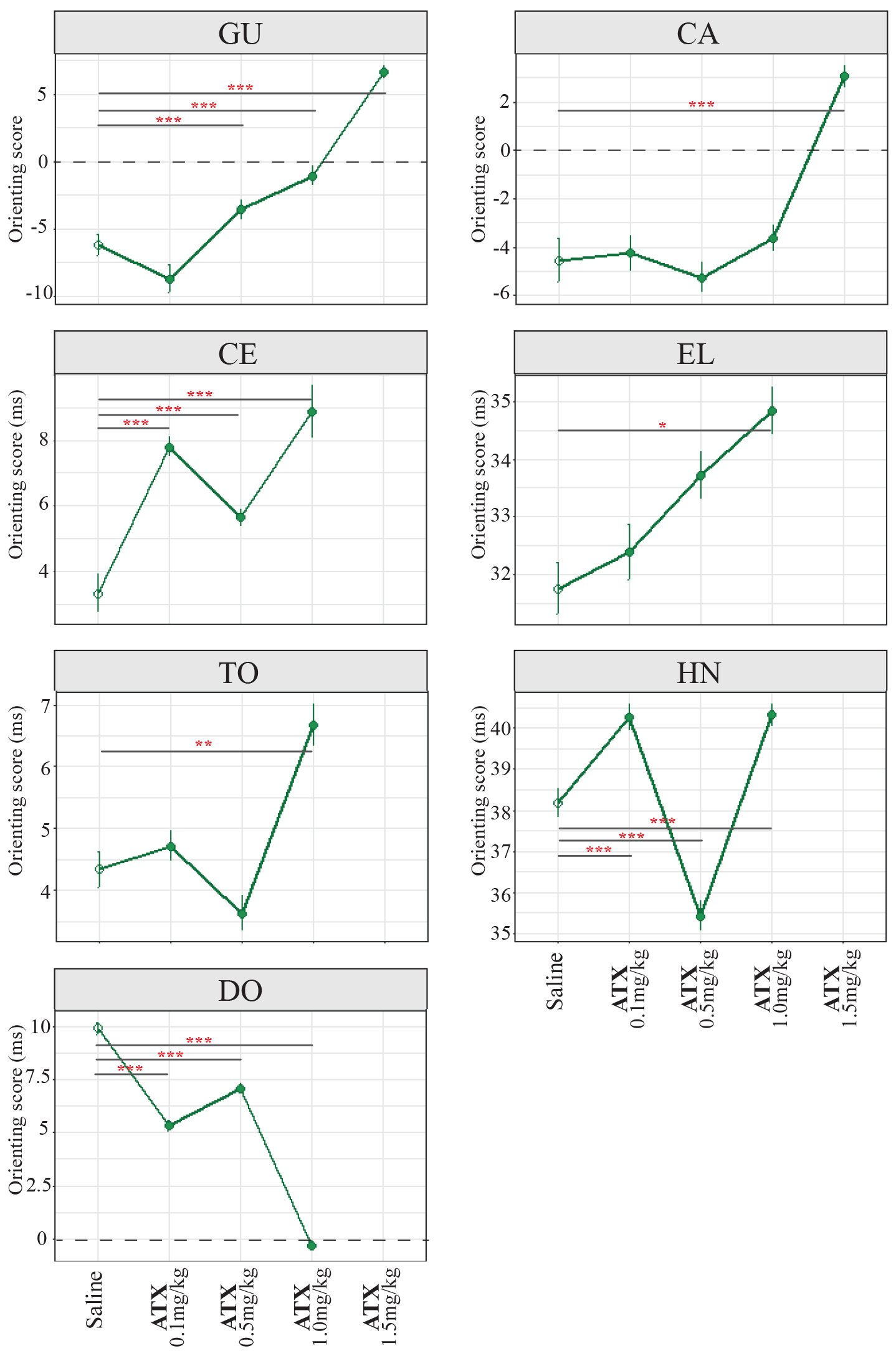
ATX effect on attentional orienting. For each animal and each pharmacological condition, we computed the normalized averaged orienting scores across runs in the different pharmacological conditions (mean±s.e). Our results show that ATX enhanced the orienting score in most monkeys. \**:p-value<0.01; **:p-value<0.05; ***:p-value<0.001*.

### 3.1. Effect of ATX in the Mixed Runs

#### 3.1.1. ATX effect on pupil size (figure 2)

We found a significant main effect of pharmacological condition on pupil diameter in all monkeys (all p-values <0.001). For all monkeys except DO, the highest dose of ATX increased the pupil diameter compared to the saline condition (all p-values <0.001). For DO, the highest dose of ATX (1.0mg/kg) significantly decreased the pupil diameter compared to the saline condition (p-value<0.001).

#### 3.1.2. ATX effect on attentional scores (Table 2)

As predicted (Posner et al., 1980), in the saline condition, 5 out of 7 monkeys exhibited a significant alerting effect, i.e. shorter RTs in neutral trials compared to non-cued trials, and a significant orienting effect, i.e. shorter RTs in valid trials compared to invalid trials (*alerting effect*: p=0.002 for CE and p<0.001 for EL, TO, HN, DO - *orienting effect*: all p’s<0.001). For all monkeys, the remote distractor led to longer RTs (all p’s<0.001) whereas the proximal distractor had different effects depending on monkeys. The proximal distractor either reduced RTs (p’s<0.001 for GU, CE and TO) or increased RTs (p’s<0.001 for EL, HN and DO).

ATX differentially modulated these attentional scores. Table 2 summarizes the effect of the highest dose of ATX (1.0mg/kg for monkeys CE, EL, TO, HN, DO and 1.5mg/kg for monkeys GU and CA). We found that attentional scores were differentially affected by ATX injection. Specifically, ATX more consistently affected the orienting process as compared to the alerting and distractor filtering processes (see also figure 3). Indeed, ATX injection significantly modified the orienting scores in all monkeys (all p’s<0.001) regardless of the pattern observed in the saline condition. Post-hoc tests revealed that ATX enhanced the orienting effect in 6 out of 7 monkeys (p<0.001 for GU, CA, CE, EL and HN; p=0.014 for TO). The enhancement of the orienting effect increased as a function of the ATX dose in 5 out of 7 monkeys (figure 3). One monkey (monkey DO) had a reversed modulation, the orienting effect decreasing as a function of the ATX dose. By comparison, our results showed that ATX either decreased or increased the alerting scores and the remote or proximal distractor scores depending on the animal.

**Table 2.**
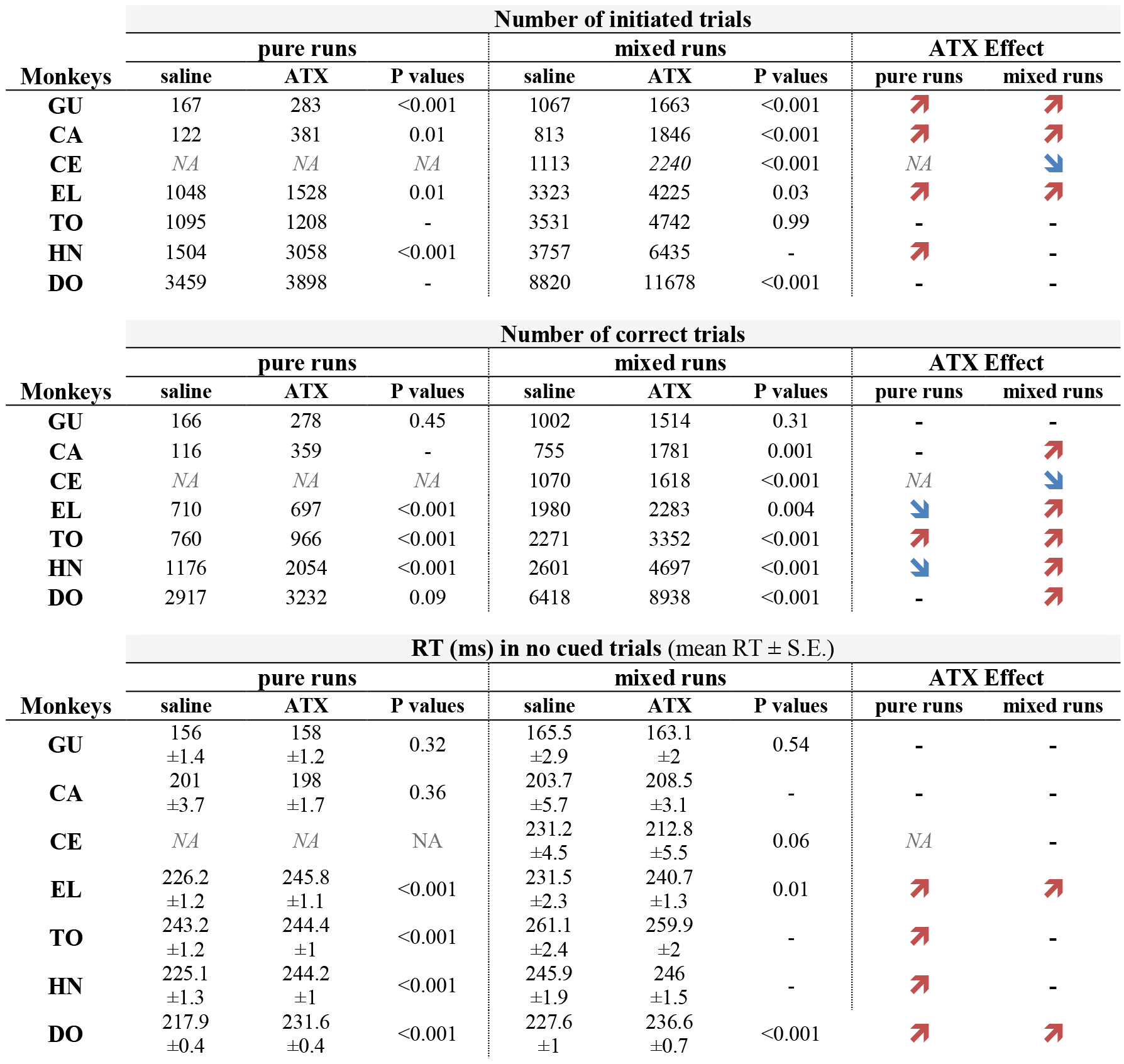
Attentional scores in mixed runs for the highest dose of ATX. (1.0mg/kg for CE, EL, TO, HN, DO and 1.5mg/kg for GU and CA). *p-values reflects pairwise comparisons between the saline and the highest dose of ATX with corrections for multiple comparisons. *: significant effect in the saline condition (p>0.05). ↗ or ↘: significant increase or decrease, respectively, after ATX administration.: -: no difference between saline and ATX conditions*. Overall, ATX modulates all attentional scores but the most consistent effect was found for the orienting score.

#### 3.1.3. ATX effect on attentional orienting

The most consistent effect of ATX on the attentional scores across animals was an improvement of the orienting effect, i.e. shorter RTs on valid than on invalid trials. To identify whether this was driven by a change in sensory accumulation or a change in decision threshold, we compare the response strategy, as assessed from RT distributions, in the saline and ATX conditions, using the LATER model. In the saline condition, in 3 out of the 5 animals exhibiting an orienting effect, this effect resulted from a lower decisional threshold in the valid compared to the invalid trials (p’s<0.001 EL, HN and DO). The two other monkeys did not exhibit any specific strategy. Under ATX, in all animals except monkey DO, the improvement of the orienting effect corresponded to a reinforcement of this decision threshold-based strategy in valid compared to invalid trials (p=0.028). For the one animal whose orienting score was significantly deteriorated under ATX (monkey DO), ATX induced the opposite effect, i.e. a switch in the strategy, from a change in the decisional threshold in the saline condition (p<0.001) toward a change of the accumulation rate in the ATX condition (p<0.001).

The enhancement of the orienting effect under ATX could result from faster RTs in both valid and invalid trials. Alternatively, it could be that ATX alters processing in only one type of trials. We thus examined the effect of ATX on the RTs in valid and invalid trials. All animals, with the exception of EL, exhibited a significant two-way interaction between pharmacological condition (Saline, ATX) and cue type (Valid, Invalid, Neutral and No cue) (GU, CA, CE, TO, HN and DO all p’s<0.001 and p=0.25 for EL). As shown in figure 4A, for the majority of monkeys, this effect was driven by shorter RTs in the valid trials (GU, CA, CE, TO, EL all p’s<0.001). RTs in the invalid trials were only marginally affected by ATX. The analysis of the RT distributions with the LATER model further demonstrated a faster accumulation rate for the valid trials in the ATX condition compared to the saline condition for 4 monkeys (EL, DO, GU and CA p’s<0.05, data not shown). For the invalid trials, ATX had no systematic impact on the RT distributions. This effect is exemplified in figure 4B for monkey EL. Overall, this indicates that the improvement of orienting induced by ATX injection is driven by faster accumulation rates following the presentation of a valid cue as compared to the saline condition.

**Figure 4.**
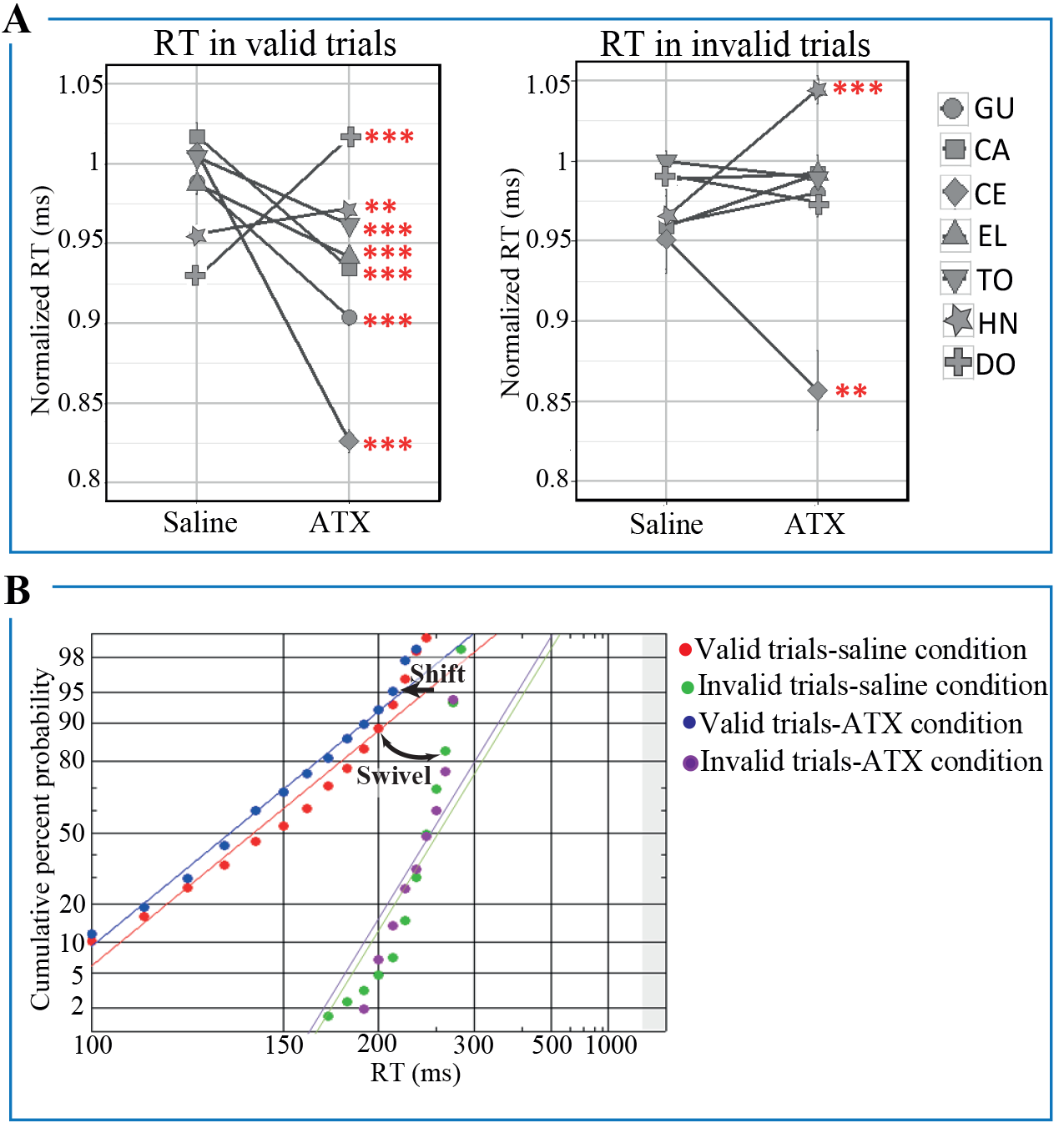
ATX effect on RTs in valid and invalid trials for the highest dose of ATX. (1.0mg/kg for CE, EL, TO, HN, DO and 1.5mg/kg for GU and CA)**. A:** For each animal and each pharmacological condition, we computed the normalized RTs in valid (left panel) and invalid (right panel) trials across runs (mean±s.e) by dividing RTs by the root mean square separately for each type of trial (valid and invalid) and each monkey. **B:** Example of reciprobit plot in valid and invalid trials in the saline and ATX conditions for monkey EL. \*\**:p-value<0.05; ***:p-value<0.001*.

**Figure 5.**
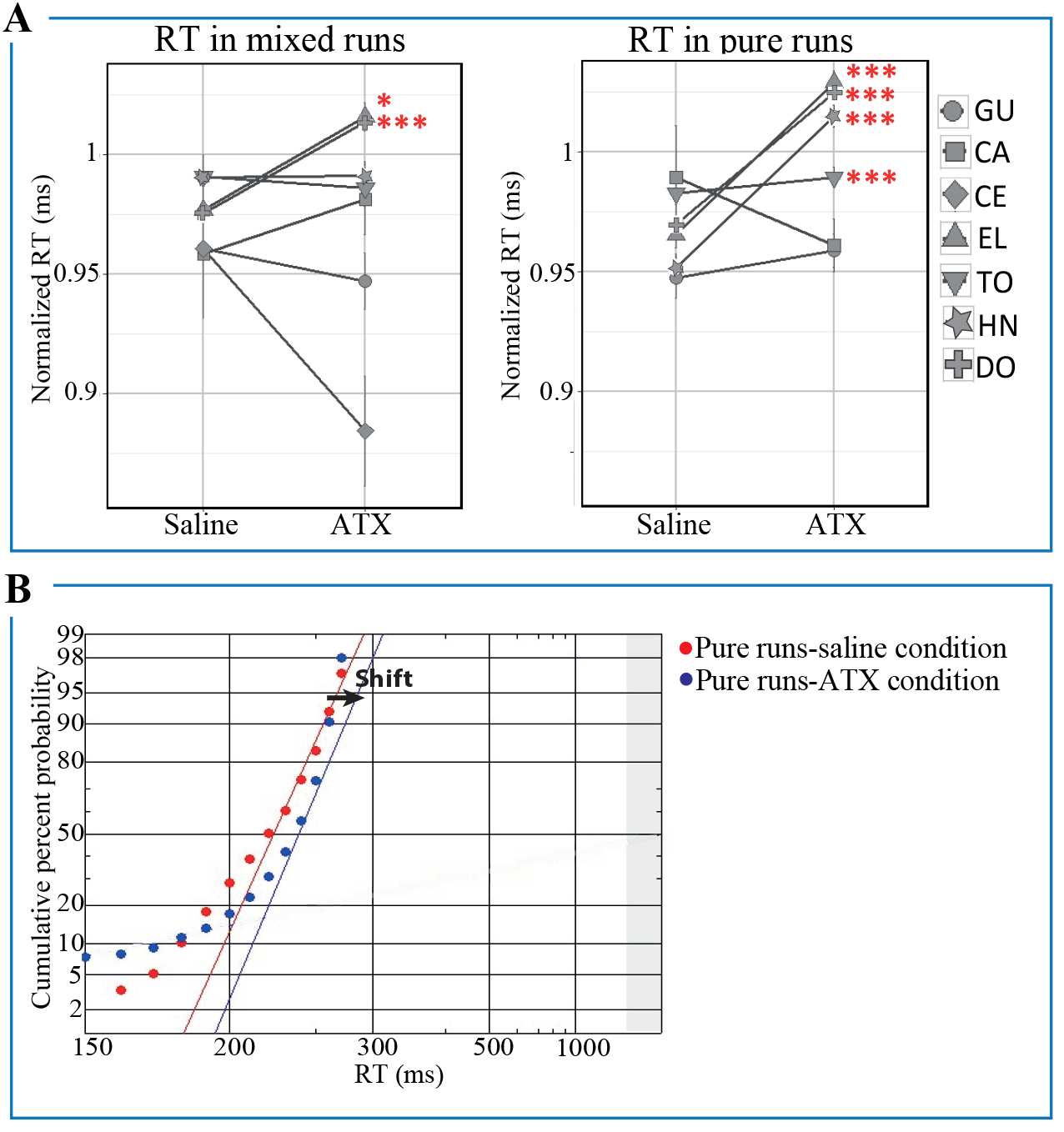
ATX effect on RTs in non-cued trials in mixed and pure runs for the highest dose of ATX. (1.0mg/kg for CE, EL, TO, HN, DO and 1.5mg/kg for GU and CA). **A:** For each animal and each pharmacological condition, we computed the normalized RTs in non-cued trials in mixed (left panel) and pure (right panel) runs across runs (mean±s.e) by dividing RTs by the root mean square separately for each type of runs (pure and mixed) and each monkey. B: Example of reciprobit plot in non-cued trials in mixed and pure runs in the saline and ATX conditions for monkey EL. \**:p-value<0.01; ***:p-value<0.001*.

### 3.2. Pure versus Mixed runs: ATX enhances task context effects.

#### 3.2.1. ATX effect on number of trials

Table 3 summarizes the effect of the highest dose of ATX (1.0mg/kg for monkeys CE, EL, TO, HN, DO and 1.5mg/kg for monkeys GU and CA) on animals’ performance in pure and mixed runs. For 3 out of 7 monkeys, ATX increased the number of initiated trials in both types of runs (mixed runs: p values < 0.001 for GU and CA; p=0.03 for EL; pure runs: p value <0.001 for GU; p=0.01 for CA and EL). By contrast, it increased the number of correct trials in only one animal (TO p value<0.001) in the pure runs compared to 5 in the mixed runs (p values <0.001 for TO, HN, DO; p=0.004 for EL). In other words, ATX equally increased the number of initiated trials in half of the animals in both mixed and pure runs while its effect on accuracy, measured as the number of correct trials, was more pronounced in the mixed runs as compared to the pure runs.

**Table 3.**
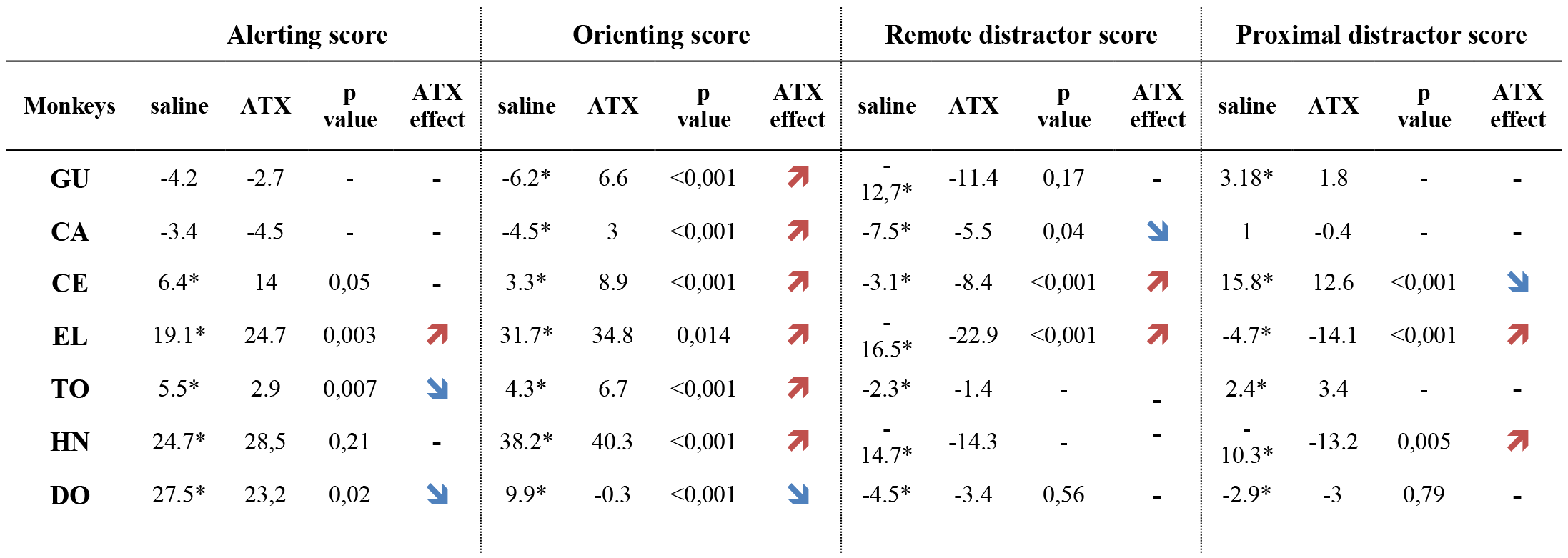
Number of trials and RTs to non-cued targets in mixed and pure runs for the highest dose of ATX. (1.0mg/kg for CE, EL, TO, HN, DO and 1.5mg/kg for GU and CA). *↗ or ↘: significant increase or decrease, respectively, after ATX administration. -: no difference between saline and ATX conditions. NA: not applicable, note that CE did not perform the pure run condition.* Overall, ATX tended to increase the number of initiated trials in both types of runs while it tended to improve accuracy only in mixed runs. In addition, ATX increased RTs in pure runs in the majority of monkeys whereas its effect on RTs was subtler in mixed runs.

#### 3.2.2. ATX effect on RTs

We then focused on RTs on the non-cued trials in the pure versus the mixed runs (Table 3). We found that ATX increased RTs in pure runs in the majority of monkeys (p values < 0.001 for EL, TO, HN, DO), whereas its effect was subtler in mixed runs where it increased RTs in only two animals (p=0.01 for EL and p value <0.001 for DO). The analysis of the RT distributions with the LATER model further demonstrated a slower accumulation rate in the pure ATX runs compared to the pure saline runs for 3 monkeys (EL and DO, p values =0.001, HN, p value <0.03) as well as compared to the mixed ATX runs for 3 monkeys (EL, HN and DO, p values <0.001).

## 4. Discussion

We tested the impact of ATX, a NE reuptake inhibitor that increases NE availability in the brain, on visuo-spatial attention, in seven monkeys performing a predictive saccadic cued task. We report two new findings. First, we found that ATX differentially impacted the three attentional scores measured in the mixed runs, namely alerting, orienting and the distractor interference effects, most consistently improving the orienting process across the animals. Second, we found that the animals were slower to detect non-cued targets, specifically in pure runs, in the ATX compared to the saline condition. Our results suggest that the NE influences specific processes of visuo-spatial attention, and that this influence depends on the context.

### 4.1. Boosting NE transmission most consistently modulates attentional orienting in a predictive context

We assessed the impact of ATX on attentional processes in mixed runs. In these runs, the cue accurately predicted the upcoming target location in 80% of the trials rendering the context highly predictive. We found that ATX affected, though not equally, all attentional processes tested in the present work, namely alerting, orienting and the distractor interference effect. Specifically, ATX changed, in a dose-dependent manner, the orienting process in all animals; deterioration did occur (1/7 monkeys), but the typical effect was an improvement (6/7 monkeys). This improvement of the orienting process resulted from faster RTs in the trials where the cue accurately predicted the location of the target (valid trials), i.e. the most prevalent trials in our task. This result is in line with two previous studies that reported that clonidine, which decreases NE transmission, attenuated the orienting process in humans (Coull et al. 2001; Clark et al. 1989) in a predictive context. In another study, Witte and Marrocco (Witte and Marrocco 1997) failed to reveal such an effect using a task in which valid trials constituted 57% of the total trials, i.e. in a task, in which the spatial cues were much less predictive than in the present study or the Coull et al. (2001) and Clark et al. (1989) studies. As a result, in the absence of a highly predictive context, monkeys probably had to rely more heavily on stimulus-driven processes as opposed to both stimulus-driven and goal-directed processes elicited by informative peripheral cues (Chica et al. 2014). This suggests that the impact of NE modulating agents might depend on the predictability of the cue and in more general terms on the context. In line with this idea, a recent study reported larger diameter of the pupil, often considered as a proxy of the LC-NE activity, in highly predictive contexts (in which the cue accurately predicted the location of the upcoming target in 80% of the trials) as compared to none predictive contexts (50%, chance level, Dragone et al. 2018). All these results suggest that the impact of NE in visuo-spatial attention might depend on the level of prediction provided by the context and might be more pronounced when attentional orienting involves highly informative and reliable cues.

In addition, our results show a different effect of ATX in pure *versus* mixed runs, the former being devoid of spatial cues and distractors as opposed to the latter one. First, ATX more consistently affected the rate of success (i.e. number of correct trials) across animals in the mixed runs compared to the pure runs. Second, when focusing on the non-cued trials in both types of runs, it appears that ATX more consistently increased RTs for these trials in pure runs while it only marginally affected RTs for these particular trials in mixed runs. In our experimental design, the monkeys performed about 3 times more mixed runs trials compared to pure runs trials. It is thus possible that the impact of ATX on performance was biased toward the most prevalent type of runs (i.e. mixed runs) and more specifically toward the most prevalent type of trials (i.e. valid trials that represented 80% of the trials, with a spatial cue accurately predicting the location of the target). At the time of testing, all the animals had extensive experience with the task and the alternations between the pure and mixed runs. We thus suggest that **t**he difference of ATX effect on pure *versus* mixed runs might be interpreted in terms of a trade-off in performance that depended on the context. This finding is in line with the idea that the LC-NE system facilitates the mobilization of sensory and attentional resources to process information of the environment (Varazzani et al. 2015) and to provide behavioral flexibility (Lapiz and Morilak 2006; Seu et al. 2009; Cain et al. 2011). NE-dependent improvement in performance has been reported in other tasks involving working memory (Gamo et al. 2010), cognitive control (Faraone et al. 2005) or sensory discrimination (Gelbard-sagiv et al. 2018). Our results further suggest that, beyond a global adjustment of the behavior to the context, ATX fine-tunes the behavior at the level of the trial to maximize reward rate, leading to a trade-off in the infrequent trials (Aston-Jones and Cohen 2005; Bouret and Sara 2005; Corbetta et al. 2008; Fazlali et al. 2016).

Thus, to answer our first question as to which components of visuo-spatial attention are under the influence of NE, our results points towards a specific effect onto the dynamic and flexible components of attention, namely spatial orienting and executive control when the context is highly predictive. Note that the effect of ATX, at the highest dose used in the present study, might have also influenced the dopamine transmission in the brain and in particular in the prefrontal cortex (Bymaster et al. 2002; Upadhyaya et al. 2013). At this stage, one cannot rule out this possibility and future studies should tackle this difficult challenge to tease apart the specificity of each of these two major neuromodulators onto attentional processes.

### 4.2. ATX-boosting effect on spatial orienting reflects changes on both sensory accumulation rate and decision threshold

The detection of a target involves both a perceptual process that can be modelled by an accumulation of information, and a decision-making step more related to top-down processes, that can be modelled by the application of a decision threshold (Noorani and Carpenter 2016). Thus, in addition to measuring the impact of a NE agent on attentional scores using median reaction times, we also sought to identify NE-driven variations in accumulation rate and decision threshold by comparing RT distributions using LATER model statistics. First, the LATER model revealed that the adaptation to the context observed under ATX condition, highlighted by a specific improvement of attentional orienting, is explained by a lower decisional threshold in ATX condition compared to saline condition. Second, we found a faster accumulation rate specifically for the trials in which the target was preceded by a predictive spatial cue (validly cued trials) under ATX with respect to saline. In other words, under high NE availability, monkeys both accumulated the available sensory evidence faster and needed less sensory information to take their decision to saccade toward the target, specifically in the prevalent valid trials. On the contrary, we observed a slower accumulation rate in the ATX condition compared to the saline condition in the pure runs. This finding is in line with an increasing number of studies showing that NE influences bottom-up processes, even at very early-stages of sensory signal processing improving the signal-noise ratio in sensory cortex in response to incoming stimuli, to shape the behavior according to the environment (see Navarra and Waterhouse 2018; Waterhouse and Navarra 2018). For example, it has been shown that following a systemic injection of ATX, neuronal responses to light stimuli was enhanced in dorsal lateral geniculate nucleus (i.e. the primary sensory relay for visual information from the retina to the visual cortex) in anesthetized rats (Navarra et al. 2013). A recent study showed that manipulating the NE level in humans modulates the perceptual sensitivity to detect a visual target and this effect reflected changes in evoked potentials and fMRI signals in visual cortex (Gelbard-Sagiv et al. 2018). At rest, ATX was also found to reduce the functional correlation strength within sensory networks and to modify the functional connectivity between the LC and the fronto-parietal attention network (Guedj et al. 2016, 2017), involved in visuo-spatial orienting (Corbetta et al. 2008)

Thus, to get to our second aim that was to characterize the specific action of NE onto the visuo-spatial components, our results points toward two complementary actions of NE, on both bottom-up and top-down processes. Our results bring new evidence to the role of NE on attentional processes. We highlight, in particular, the impact of the context (predictive *versus* non-predictive) on its effect on attentional processes. We also pinpoint its complex mechanism of action on spatial attention, exerted at different levels, likely reflecting changes within sensory cortex leading to faster accumulation rate to incoming stimuli as well as the adjustment of the decisional threshold via an action of NE within prefrontal regions (Robbins and Arnsten 2009; Arnsten 2011; Arnsten and Pliszka 2011).

## Acknowledgements

We thank Gislène Gardechaux, Frédéric Volland, Roméo Salemme, Eric Koun, Jean-Luc Charieau, Fidji Francioly and Serge Pinède for technical and engineering assistance. This work was funded by the French National Research Agency (ANR) ANR-14-CE13-0005-1 grant. It was also supported by the NEURODIS Foundation and the James S. McDonnell Scholar award. It was performed within the framework of the LABEX CORTEX (ANR-11-LABX-0042) of Lyon University within the program “Investissements d’Avenir” (ANR-11-IDEX-0007) operated by the ANR.

## Conflict of Interest

The authors declare no competing financial interests.

